# Image-based Motion Artifact Reduction on Liver Dynamic Contrast Enhanced MRI

**DOI:** 10.1101/2021.04.23.441167

**Authors:** Yunan Wu, Junchi Liu, Gregory M White, Jie Deng

**Affiliations:** Department of Diagnostic Radiology, Rush University Medical Center, 1653 W. Congress Pkwy, Jelke Ste 181, Chicago, IL, USA 60612; Department of Electrical Computer Engineering, Northwestern University, 633 Clark Street, Evanston, IL, USA 60208; Medical Imaging Research Center and Department of Electrical and Computer Engineering, Illinois Institute of Technology, Chicago, IL, USA 60616

**Keywords:** motion artifacts, liver, dynamic contrast enhanced MRI, deep learning, simulation, perceptual loss

## Abstract

Liver MRI images often suffer degraded quality from ghosting or blurring artifact caused by patient respiratory or bulk motion. In this study, we developed a two-stage deep learning model to reduce motion artifact on dynamic contrast enhanced (DCE) liver MRIs. The stage-I network utilized a deep residual network with a densely connected multi-resolution block (DRN-DCMB) network to remove the majority of motion artifacts. The stage-II network applied the perceptual loss to preserve image structural features by updating the parameters of the stage-I network via backpropagation. The stage-I network was trained using small image patches simulated with five types of motion, i.e., rotational, sinusoidal, random, elastic deformation and through-plane, to mimic actual liver motion patterns. The stage-II network training used full-size images with the same types of motion as the stage-I network. The motion reduction deep learning model was testing using simulated motion images and images with real motion artifacts. The resulted images after two-stage processing demonstrated substantially reduced motion artifacts while preserved anatomic details without image blurriness. This model outperformed existing methods of motion reduction artifact on liver DCE-MRI.

## 1. Introduction

Magnetic resonance imaging (MRI) is the most commonly used imaging modality for diagnosis of liver cancer and other liver diseases. Dynamic contrast enhanced (DCE) MRI is an essential sequence in liver MRI protocol to evaluate liver tissue perfusion changes (1) for the assessment of liver fibrosis and cirrhosis (2,3), and the differentiation of benign from malignant tumors (4). The liver DCE-MRI sequence is continuously acquired before and after the intravenous administration of gadolinium-based contrast agent at several time points and requires patients to hold their breath for 15-25 seconds during each imaging acquisition. The critical timing required for this sequence allows for only a few seconds of patient breathing time between imaging acquisitions which causes patient fatigue and thus respiratory motion artifacts which degrade image quality through ghosting artifact or blurring of the images (5).

In clinical MRI examinations, patients are given written and visual instructions to practice breath-holding required for imaging which can help reduce the number of repeat imaging (6) acquisitions to correct artifacts. However, once the contrast agent is injected, it is not practical to stop and repeat the DCE image acquisition if patient respiratory motion occurs because of the timing requirements and gadolinium toxicity. Advanced image reconstruction techniques such as compressed sensing (CS) have been used to remove respiratory motion artifacts (7–10), where signals are reconstructed from the highly under-sampled signal acquisitions to produce an MR image within a shortened scan time. Radial K-space sampling combined with CS and parallel imaging enabled free breathing acquisitions of liver DCE-MRI (9). Although the CS has been proved to help motion reduction, it was limited by the acceleration rate and availability from different MRI manufacturers and platforms. Also, patients with irregular breathing patterns are not ideal for use with the CS technique. Therefore, the development of image-based post-processing methods on the DCE images will be useful to retrospectively mitigate motion artifacts and improve image quality.

Recently, deep learning (DL) approaches have been developed in medical imaging use cases including image reconstruction and artifact reduction (11,12), motion detection and correction (13), and image quality control (14,15). DL methods utilized convolutional neural networks (CNNs) to extract features of different types of artifacts and correct them in brain (16,17), abdominal (18–20) and cardiac imaging (13). Specifically for liver DCE-MRI, Tamada et al. proposed a denoising CNN on multi-phase magnitude-only image patches that learned the artifact patterns as residual feature maps and then subtracted them from the original images to obtain the motion reduced images (19). Kromrey et al. developed deep learning filters in CNNs for multi-arterial phase acquisitions and improved image quality on severely degraded images (20). However, the motion artifacts used for model training in both studies were simulated based on specific re-ordering of K-space lines, which cannot represent the full spectrum of motion artifact patterns. It is necessary to create a more generalized CNN model that is able to reduce various degrees and types of motion artifacts occurring on liver DCE-MRI.

In this study, we developed a two-stage DL model to reduce motion artifacts on liver DCE-MRI. Stage-I utilized a deep residual network with densely connected multi-resolution blocks (DRN-DCMB) to remove the major artifacts of the image (16). Stage-II exploited the perceptual similarity in a high-dimensional feature space exacted by a VGG-16 network to preserve imaging details (21). The combination of these two networks realized removal of motion artifacts while preventing the over-smoothing effects. To increase the generalizability of this model in more complicated situations, motion artifact patterns induced by various types and degrees of motion were simulated during the training process. Finally, we tested whether this proposed DL method may effectively reduce the real motion artifacts occurring on liver DCE-MRI images.

## 2. Materials and Methods

### 2.1 Liver MRI Dataset

This retrospective study was approved by the Rush University Medical Center institutional review board and written informed consent was waived. A dynamic multi-phase DCE imaging protocol covering the whole liver volume was acquired before and after intravenous (IV) contrast administration (0.1 mL/kg Eovist^®^ Gadoxetate Disodium) with an injection rate of 2 mL/sec. Imaging parameters for each acquisition were: DCE imaging was acquired using the three-dimensional (3D) T1-weighted (T1W) spoiled gradient echo (SPGR) sequence on different 1.5T/3.0T MRI scanners (Signa Artist, MR450W, and Excite manufactured by GE Healthcare, Magnetom Espress and Verio manufactured by Siemens Healthineers) with breath-hold time range of 15-25 seconds. Axial stacks of DCE images were acquired before and after IV contrast injection at early arterial, late arterial, portal venous, and transitional phases, followed by the hepatobiliary phase 20 minutes after injection. Pre- and post-contrast DCE images were mixed for model training and testing purposes.

### 2.2 Motion Artifact Simulation

For model training process, pairs of images without motion (i.e. clean images) and with simulated motion artifacts (i.e. simulated motion images) were generated by manipulating the K-space data of the clean images and/or transformed images. The signal phases or the order of a certain range of K-space data were altered to simulate different types and degrees of motion artifacts (22,23). The simulation steps were described in **Fig. 1**. (i) A given image, which can be either a clean image or a transformed image, was converted to its K-space data by Fast Fourier Transform (FFT); (ii) K-space data of the clean image and/or transformed image was manipulated according to different rules described in section 2.2.1-2.2.5 to form a new K-space. (iii) A simulated motion image was reconstructed from the new K-space using inversed FFT (iFFT). The simulation process can be generalized in Eq. 1,

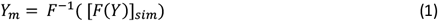

**Figure 1.**
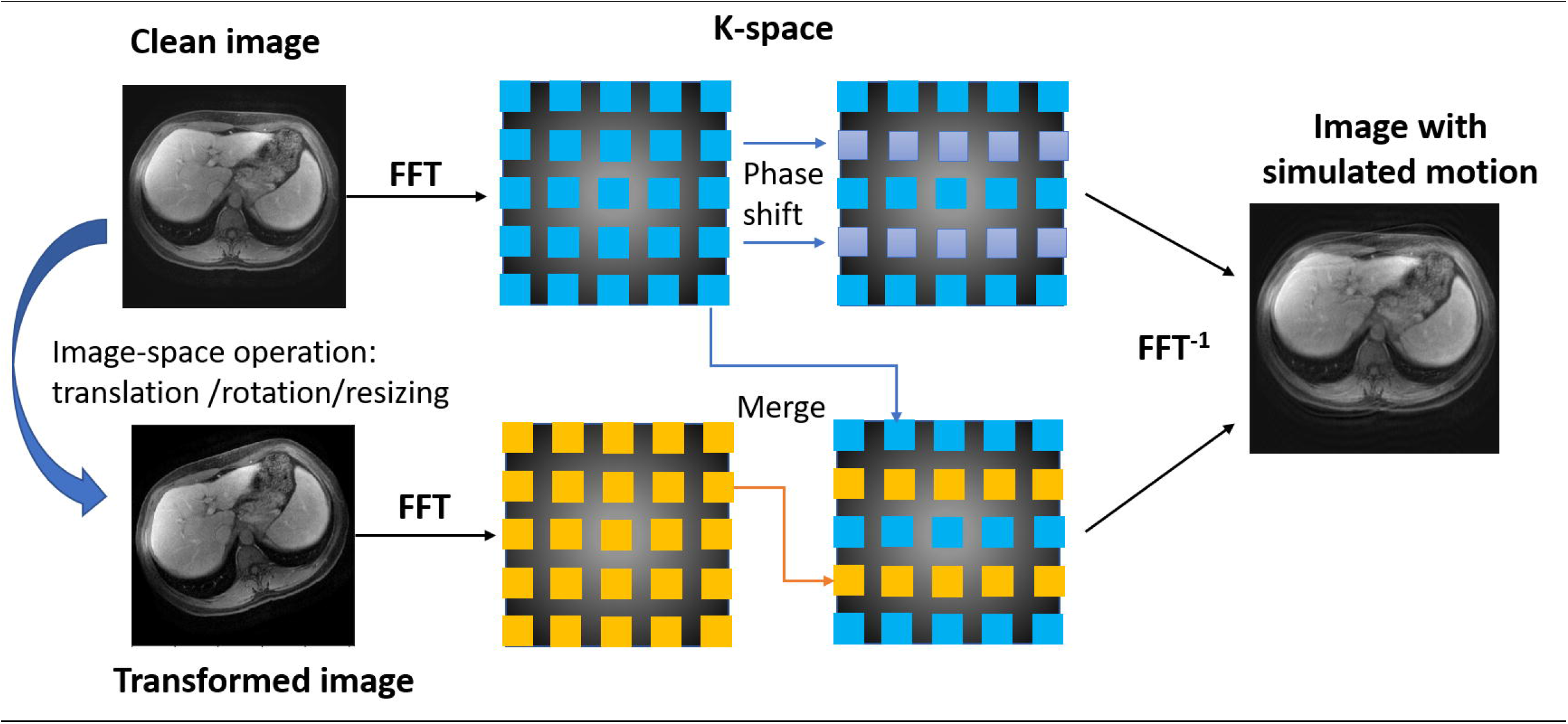
MRI motion artifact simulation process. The K-space data of a clean image and/or the corresponding transformed image was manipulated to generate a new K-space with simulated motion, which went through iFFT to form a new image with simulated motion artifacts. The pairs of the clean image and new images with different types of simulated motion artifacts were used for model training.

where *F* and *F*^−1^ denote FFT and iFFT, *Y* and *Y*_*m*_ represent the clean image and the image with simulated motion artifacts, and [·]_*sim*_ is the simulated K-space with motion.

Five different types of motion were simulated to mimic realistic motion artifacts (section 2.2.1-2.2.5), and the corresponding residual artifact images between the clean images and simulated motion images were shown in **Fig. 2**.

**Figure 2.**
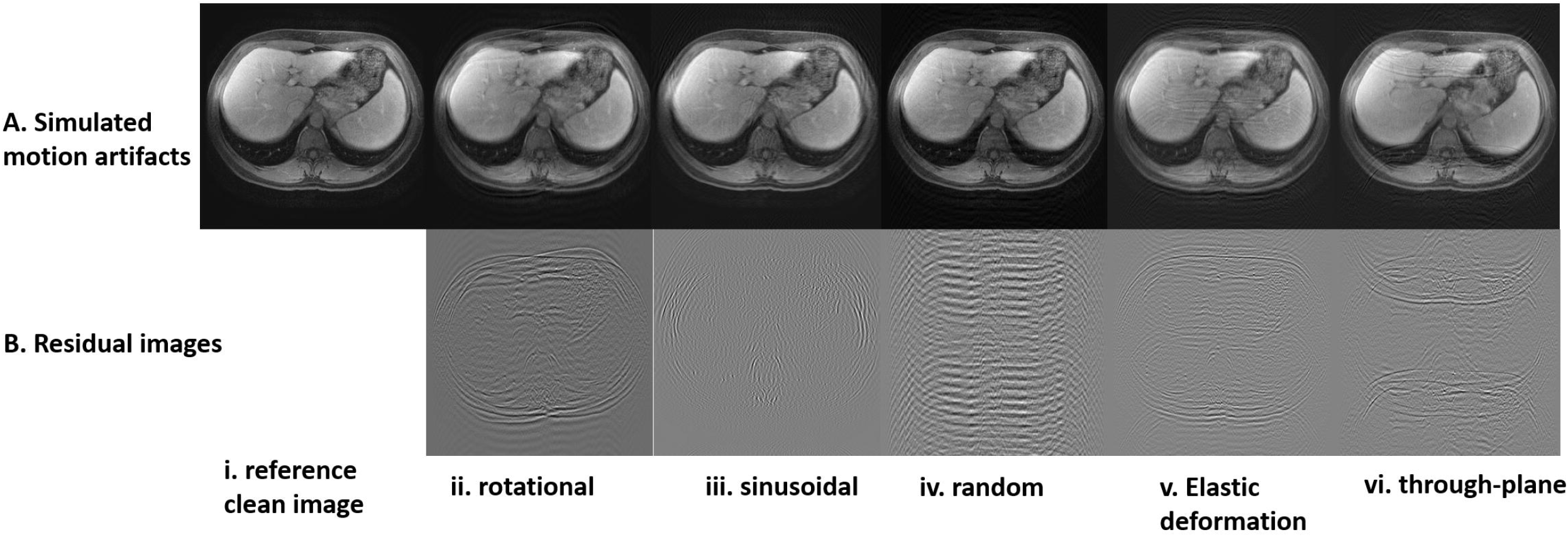
Examples of a clean image (i) and corresponding images bearing five different types of simulated motion artifacts (ii-vi) (row A). The residual images represented the motion patterns of each simulated image by subtracting the reference clean image (row B).

#### 2.2.1 Type 1: Rotational Motion

One of the major motion artifacts is bulk body movement, which causes incoherent ghosting and blurring on images (23). The clean image was rotated to create a rotated image with a random rotational angle between -20° and 20°, which then went through FFT to form the rotated K-space *F*(*Y*_*r*_). A new K-space [*F*(*Y*)]_*rotation*_ (i.e., [*F*(*Y*)]_*sim*_ in Eq.1) was generated as follows:

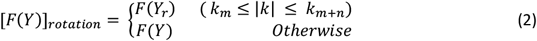

where *F*(*Y*_*r*_) represents the K-space domain of the rotated image, *k* is the K-space coordinate in the phase-encoding direction (−*π* < *k* < *π*), *n* is the number of continuous K-space lines (0< *n* < 128) in *F*(*Y*_*r*_), and *m* is an index of K-space location (0 < *m* < 384). In each simulated scenario, a random number of *n* K-space lines from *F*(*Y*_*r*_), starting at a randomly chosen K-space index *m*, were filled into the new K-space [*F*(*Y*)]_*rotation*_, and the rest of the original K-space *F*(*Y*) lines were kept in [*F*(*Y*)]_*rotation*_.

#### 2.2.2 Type 2: Sinusoidal Motion

To simulate periodic human respiratory cycles, sinusoidal motion patterns were generated by changing the duration, frequency and phase of the simulated sinusoidal wave (16,19,24). The new k-space [*F*(*Y*)]_*sin*_ (i.e., [*F*(*Y*)]_*sim*_ in Eq.1) was generated by altering the signal phase of the original K-space *F*(*Y*), defined as follows:

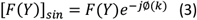

where ∅(*k*) denotes the phase shift error added to a given K-space line *k* along the phase-encoding direction, and ∅(*k*) is defined as:

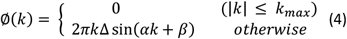

where *k*_*max*_ is the range of center K-space lines that were preserved without adding phase shift errors, and *k*_*max*_ was randomly chosen from *π*/10 to *π*/2. Δ is the number of pixels (0 < Δ < 20), depicting the severity of motion, α is the frequency of the respiratory cycle (0.1< α <5 Hz), and *β* is the phase of the respiratory wave (0 < *β* < *π*/4). Phase shift defined by ∅(*k*) was added to each K-space line *k* that was located outside the center range of (−*k*_*max*_, *k*_*max*_).

#### 2.2.3 Type 3: Random Motion

To simulate the irregular non-periodic respiratory motion, a new K-space with random motion [*F*(*Y*)]_*rand*_ (i.e., [*F*(*Y*)]_*sim*_ in Eq.1) was generated by adding signal phase shifts ∅(*k*)_*rand*_ to 10%-50% randomly selected peripheral K-space lines, whereas the center 4% - 10% of the original K-space *F*(*Y*) lines were kept intact (25). Different percentage of the preserved center K-space lines represented different levels of motion severity (24).

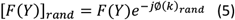

#### 2.2.4 Type 4: Elastic Deformation

Elastic deformation is often observed in abdominal MRI images and typically consists of stretching and shearing along different directions (23). Three types of motion (translation, rotation, and resizing) were combined to mimic image deformation. A clean image was shifted up or down by 10 pixels to form a translated image *Y*_*trans*_. A clean image was rotated by [-20°, 20°] to form a rotated image *Y*_*rot*_. A clean image was resized and interpolated along x and y directions to form a stretched image *Y*_*resize*_. After that, three corresponding K-space *F*(*Y*_*trans*_), *F*(*Y*_*rot*_) and *F*(*Y*_*resize*_) were obtained through FFT. Finally, a new K-space [*F*(*Y*)]_*deform*_ (i.e., [*F*(*Y*)]_*sim*_ in Eq.1) was created by interleaving randomly-selected K-space lines from *F*(*Y*_*trans*_), *F*(*Y*_*rot*_) and *F*(*Y*_*resize*_), while keeping the center 4% - 10% of the original K-space *F*(*Y*) lines of the clean image.

#### 2.2.5 Type 5: Through-plane motion

Sudden patient body position changes including movement from insufficient breath hold, lead to motion along the head-to-foot direction (23). For example, suppose slice *Y*_*s*_ is the target axial slice, slice *Y*_*s*+2_ and slice *Y*_*s*−2_ are two axial slices above the below *Y*_*s*_. These three clean images were converted to the corresponding K-space *F*(*Y*_*s*+2_), *F*(*Y*_*s*_) and *F*(*Y*_*s*−2_), respectively. A new K-space with through-plane motion [*F*(*Y*)]_*thru*_ (i.e., [*F*(*Y*)]_*sim*_ in Eq.1) was created by interleaving peripheral K-space lines randomly selected from *F*(_*s*+2_), *F*(*Y*_*s*_) and *F*(*Y*_*s*−2_), while keeping the center 4% - 10% of the original K-space *F*(*Y*_*s*_) lines.

### 2.3 Model Architecture

In this study, a two-stage deep CNN model was developed to reduce motion artifacts of the liver DCE-MRI images. The stage-I model was adapted from the DRN-DCMB network originally proposed by Liu et al. to reduce the major patterns of motion artifacts (16) (**Fig. 3**). It has three major components: multi-resolution block with U-net as inner structure, dense connection among multi-resolution blocks, and residual learning. Stage-I model was trained independently with the pairs of clean and simulated motion image patches. Next, Stage-II model utilized a “J”-shaped architecture (J-net) to optimize Stage-I model by comparing the perceptual similarity between the ground truth clean image and the output “artifact-reduced” image from the stage-I model. The perceptual loss was used to update the parameters in the stage-I model to restore more image details (26) (**Fig. 4**). The purpose of the two-stage model was to reduce the motion artifact while preserving the sharpness and resolution of the image.

**Figure 3.**
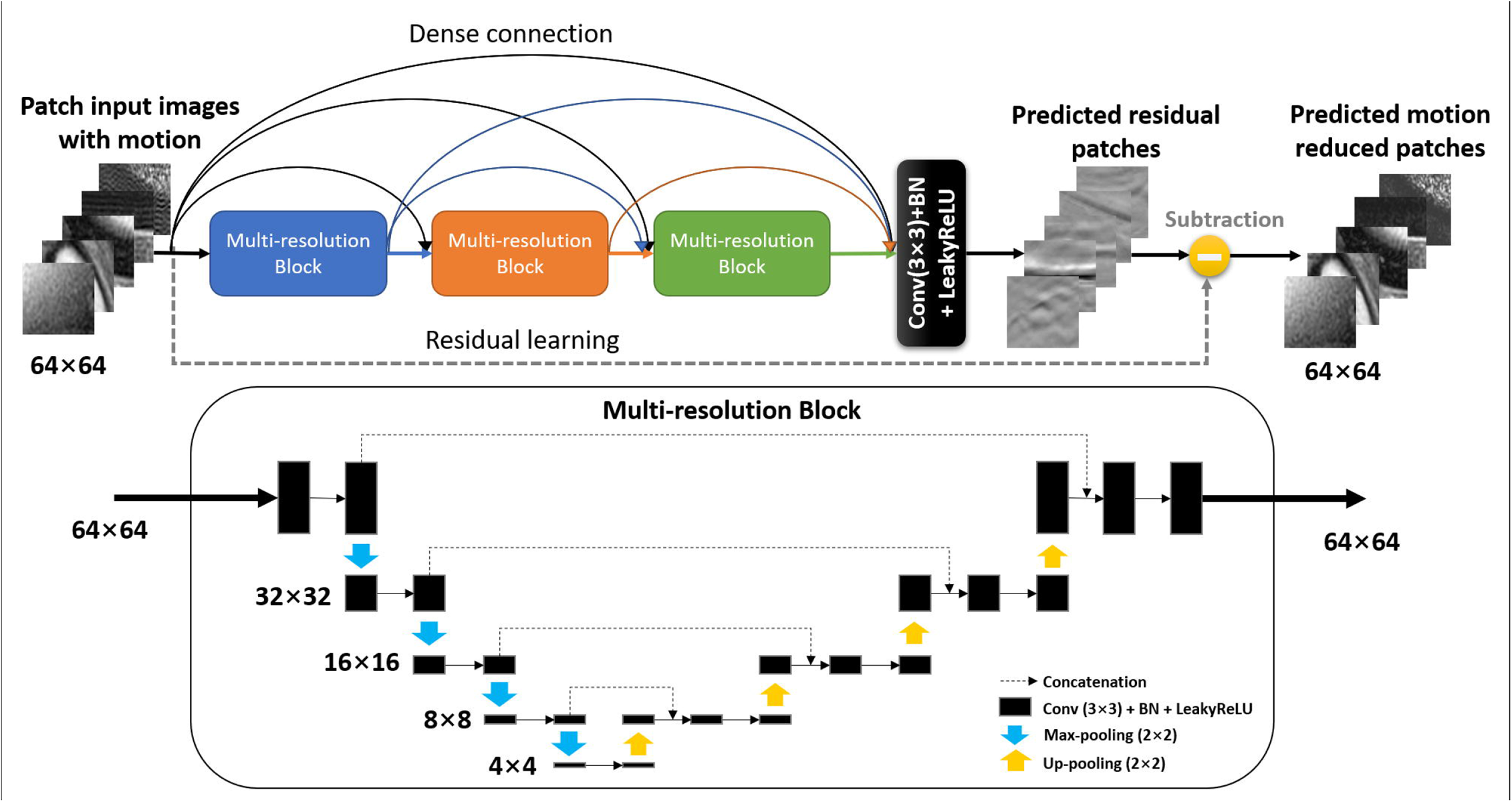
The architecture of the DRN-DCMB network (Stage-I model). Each multi-resolution block was built upon the U-net architecture.

**Figure 4.**
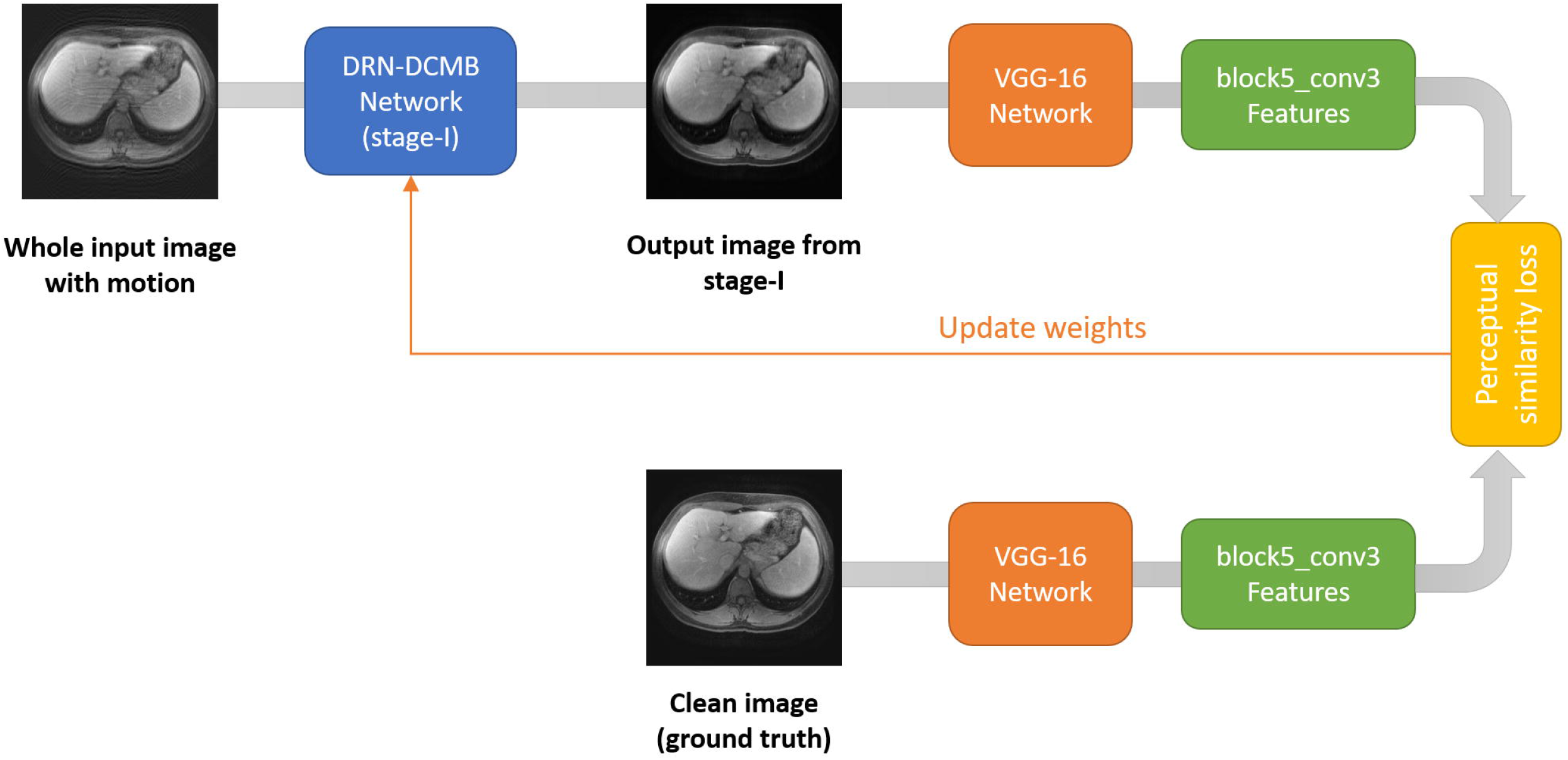
The architecture of the J-net model (Stage-II model). Perceptual loss was calculated between the feature maps extracted from the output image of the stage-I model on the long end and those extracted from the ground truth clean image on the short end of the J-net. Perceptual loss was used to further update the weights of the stage-I model.

#### 2.3.1 Stage-I: DRN-DCMB Network

As shown in **Fig 3**, the architecture of the deep residual network with densely connected multi-resolution blocks (DRN-DCMB) was composed of 3 multi-resolution blocks, followed by a basic convolutional block, which included a 3×3 convolutional layer, a batch normalization (BN) layer and a Leaky rectified linear unit (LeakyReLU) activation function. The BN layers were used to promote faster training and make the nonlinear activation functions viable (27). LeakyReLU was chosen due to its small slope for negative values in order to fix the “dying ReLU” problem and speed up the training (28). The U-net inner structure of each multi-resolution block consisted of a down-sampling path and an up-sampling path. Four 2×2 max-pooling layers were used in the down-sampling path to generate deeper levels of feature maps for extraction of local image details, such as vessels and tumors in the liver. Likewise, four 2×2 up-pooling layers were used to restore the features maps back to same size of the input image at each level. In particular, the feature maps from the down-sampling layers were concatenated to their corresponding up-sampling layers. By doing so, the global information such as organs and motion artifacts extracted by the down-sampling levels were combined with the local information exacted on the same levels along the up-sampling path. The 3 multi-resolution blocks were densely connected, in which the output from one previous multi-resolution block was concatenated to the inputs of all subsequent multi-resolution blocks. Dense connection between multi-resolution blocks preserved important learning features and therefore accelerated training speed without relearning redundant features. In addition, residual learning (29) was used during the training process to learn the residual differences (i.e. artifacts) between the simulated motion image and the clean image. The model output image was generated by subtracting the residual map from the input image. Lastly, the fully convolutional layers in the DRN-DCMB model allowed the input image size to vary, for example, small image patches (64×64) randomly sampled from the full-size image were used as the input for training purpose, and the full-size images (512×512) were used as the input during the testing process. All trainable parameters were updated by minimizing the pixel-by-pixel mean square error (MSE) loss between the clean image and the DRN-DCMB model output image. However, using the pixel-to-pixel MSE loss may cause blurring and over-smoothness problem (26,30), also observed in this study. Therefore, a stage-II model was developed to further optimize the parameters (i.e. weights) in the stage-I model to mitigate image blurriness and restore image details.

#### 2.3.2 Stage-II: J-net Network

At stage-II, the J-net exploited the perceptual similarity measurement as the loss function (31) to update the weights of the stage-I model. The perceptual loss aimed to minimize the differences in high-dimensional features between the ground truth clean image and the output image of the stage-I model. Let *M*_*i*_(·) be the feature map generated from a given layer *i*, and the feature map has the size of *h* × *w* × *c*, where h and *w* represents the height and width of the feature map, and *c* represents the number of channels. Then, the perceptual loss 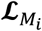 was defined as:

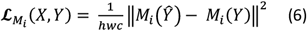

where *Ŷ* denotes the output image from the stage-I model on the long end of the J-net and *Y* denotes the ground truth clean image on the short end of the J-net (**Fig. 4**).

The initialized weights of the J-net consisted of two parts: weights trained from the stage-I model and weights of the original VGG-16 net. The VGG-16 network (32) used in J-net (21) was pre-trained on the ImageNet (33) to extract meaningful features more easily recognized by human eyes. The VGG-16 network extracted feature maps from stage-I model output image and the ground truth clean image separately, and then the perceptual loss was calculated from these two features maps, which was then back-propagated to update the weights in the stage-I DRN-DCMB network. Note all weights of the VGG-16 net itself was not trained or updated. More specifically, the stage-II model training process included the following steps. 1) The input simulated motion images to the DRN-DCMB network was a full-size image instead of small-patch images; 2) The output image from the DRN-DCMB network was fed into the pre-trained VGG-16 network for feature extraction from the layer of “block5_conv3” (feature map size: 32×32×512); 3) The ground true clean image was also fed into another pre-trained VGG-16 network for feature extraction from the layer of “block5_conv3” (feature map size: 32×32×512); 4) The perceptual loss was calculated by comparing the two feature maps, as illustrated in Eq. 5; 5) The perceptual loss was back-propagated to update the weights in the DRN-DCMB model.

### 2.4 Model Training, Validation and Testing

The liver DCE-MRI image volumes that had no obvious motion artifacts (i.e. “clean image”) in 10 patients were selected in the training process, of which 8 was used for training and 2 for validation. Five different types of the simulated motion artifacts were added to these clean images (see section 2.2). During the stage-I training, 20 small image patches (size: 64×64) were randomly generated from each full-size image, leading to a total of 200,000 patches (20 patches × 25 slices × 5 motion types × 10 randomized parameters × 8 patients) for the training dataset, and 50,000 patches (20 patches × 25 slices ×5 motion types × 10 randomized parameters × 2 patients) for the validation dataset. Other training parameters included: batch size = 64, early stopping at the 34^th^ epoch, and a learning rate initialized from 0.0001 using the Adam optimization algorithm (34). The training time was 3.7 hours for the stage-I model.

During the stage-II training process, the full-size simulated motion images (size: 512×512) were used as the input, with 10,000 images (25 slices ×5 motion types x 10 randomized parameters ×8 patients) used for training and 2500 images (25 slices ×5 motion types × 10 randomized parameters × 2 patients) used for validation. Other training parameters included: batch size = 8, early stopping at the 9^th^ epoch, and a learning rate initialized from 0.00005 using the Adam optimization algorithm. The training time was 2.6 hours for the stage-II model Two testing datasets were used to evaluate the model performance. Testing dataset-I consisted of 6250 clean images with simulated motion acquired from 5 patients (25 slices ×5 motion types × 10 randomized parameters × 5 patients). Testing dataset-II consisted of 28 images that demonstrated obvious motion artifacts acquired from another 12 patients.

All training and testing processes were performed using Tensorflow2.0 in Python3.7 on a single GPU (NVIDIA GeForce RTX 2070 Super) workstation.

### 2.5 Model Performance Evaluation

MSE and structural similarity index (SSIM) (35) were measured in the training dataset and the testing dataset-I with simulated motion. The mean squared difference between the clean image and the model output motion-reduced image was measured by MSE. SSIM is a widely used perceptual metric that addresses the differences in the structural, luminance, and contrast between two images (Eq. 7).

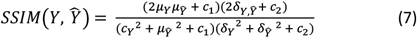

where *Y* and *Ŷ* denotes the clean and model output motion-reduced image, respectively; μ is the mean intensity and δ is the standard deviation of an image; *c*_1_ = 0.01 and *c*_2_ = 0.03 are the constants.

### 2.6 Statistical Analysis

MSE and SSIM values were calculated for each type and across all types of the simulated motion artifacts in training dataset and testing dataset-I. The SSIM and MSE values were compared en the output images generated by stage-I and stage-II models using the pair *t*-test. A p value < 0.05 indicated significant differences in comparison. All statistical analyses were performed using the SciPy library in Python 3.7.

## 3. Results

Among the 6250 testing images with simulated motion artifacts, the performance of artifact reduction by using the stage-I DRN-DCMB model alone and using the combined stage-I and stage-II (J-net) models was compared, as shown in **Table 1**. The overall SSIM of using the stage-I model was 0.947 ± 0.016 and MSE was 5.8 ± 4.0 × 10^−4^. The stage-I model achieved the best performance for the random motion simulation (SSIM: 0.956 ± 0.005, MSE: 5.2 ± 2.0 × 10^−4^). The J-net model improved the overall performance (SSIM: 0.974 ± 0.016, MSE: 4.7 ± 3.8 × 10^−4^) compared with the stage-I model alone with statistical significance (*p* < 0.05). The J-net model also achieved the best performance for random motion simulation (SSIM: 0.981 ± 0.004, MSE: 4.6 ± 1.8 × e^-4^). The relatively large standard deviation of MSE was due to the large inter-subject variations of motion artifact simulation added on the image. The performances of all types of motion were significantly improved by using the J-net model compared with the stage-I model alone (p < 0.05).

**Table 1.** Model performance comparisons between the stage-I and stage-II models using the metrics of structural similarity index (SSIM) and mean square error (MSE) for each type and over all types of simulated motion artifacts. *indicated statistical significance (p< 0.05).

| Metrics<br>Type<br>of Artifact | Stage-II |  | Stage-I |  |
| --- | --- | --- | --- | --- |
|  | SSIM | MSE | SSIM | MSE |
| Rotational | $0.978 \pm 0.022^*$ | $0.00047 \pm 0.00020^*$ | $0.951 \pm 0.024$ | $0.00054 \pm 0.00028$ |
| Sinusoidal | $0.965 \pm 0.016^*$ | $0.00053 \pm 0.00020^*$ | $0.937 \pm 0.019$ | $0.00060 \pm 0.00035$ |
| Random | $0.981 \pm 0.004^*$ | $0.00046 \pm 0.00018^*$ | $0.956 \pm 0.005$ | $0.00052 \pm 0.00020$ |
| Elastic | $0.971 \pm 0.014^*$ | $0.00046 \pm 0.00018^*$ | $0.943 \pm 0.015$ | $0.00052 \pm 0.00018$ |
| Through-plane | $0.971 \pm 0.006^*$ | $0.00027 \pm 0.00009^*$ | $0.938 \pm 0.017$ | $0.00041 \pm 0.00019$ |
| **Overall** | **$0.974 \pm 0.016^*$** | **$0.00047 \pm 0.00038^*$** | **$0.947 \pm 0.016$** | **$0.00058 \pm 0.00041$** |

\* Statistically significant ( $p < 0.05$ )

Four representative images with simulated motion artifacts and their corresponding artifact-reduced images processed by the stage-I model alone and by the J-net model were shown in **Fig. 5**. Motion artifacts were dramatically reduced in the output images generated by the stage-I model alone, however, image blurriness was also observed (**Fig. 5**, column ii). In contrast, the J-net model (**Fig. 5**, column iii) not only mitigated the motion artifacts but also preserve the image sharpness of the original clean images (**Fig. 5**, column iv).

**Figure 5.**
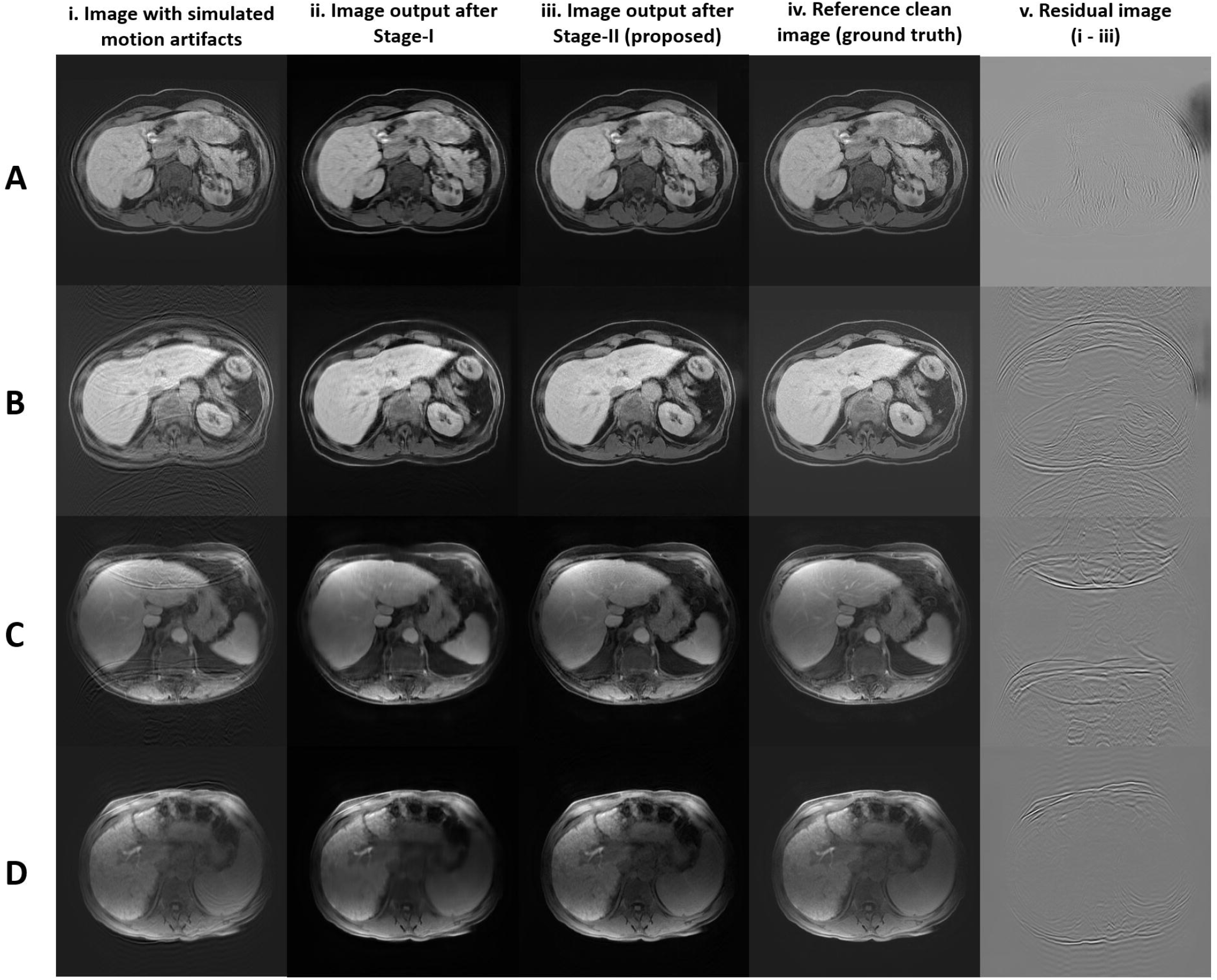
Comparisons of artifact reduction after the stage-I model (column ii) and stage-II model (column iii) on four testing motion images (column i) generated by adding motion artifacts to the reference clean images (column iv). The residual images (column v) were generated by subtracting the final motion-reduced images (column iii) from the testing motion image (column i).

Three representative images with real motion artifacts with their corresponding artifact-reduced images processed by the J-net model, and the residual artifact images were shown in **Fig. 6**. Obvious motion artifact reduction, especially in regions with severe ghosting artifacts, was observed on these images.

**Figure 6.**
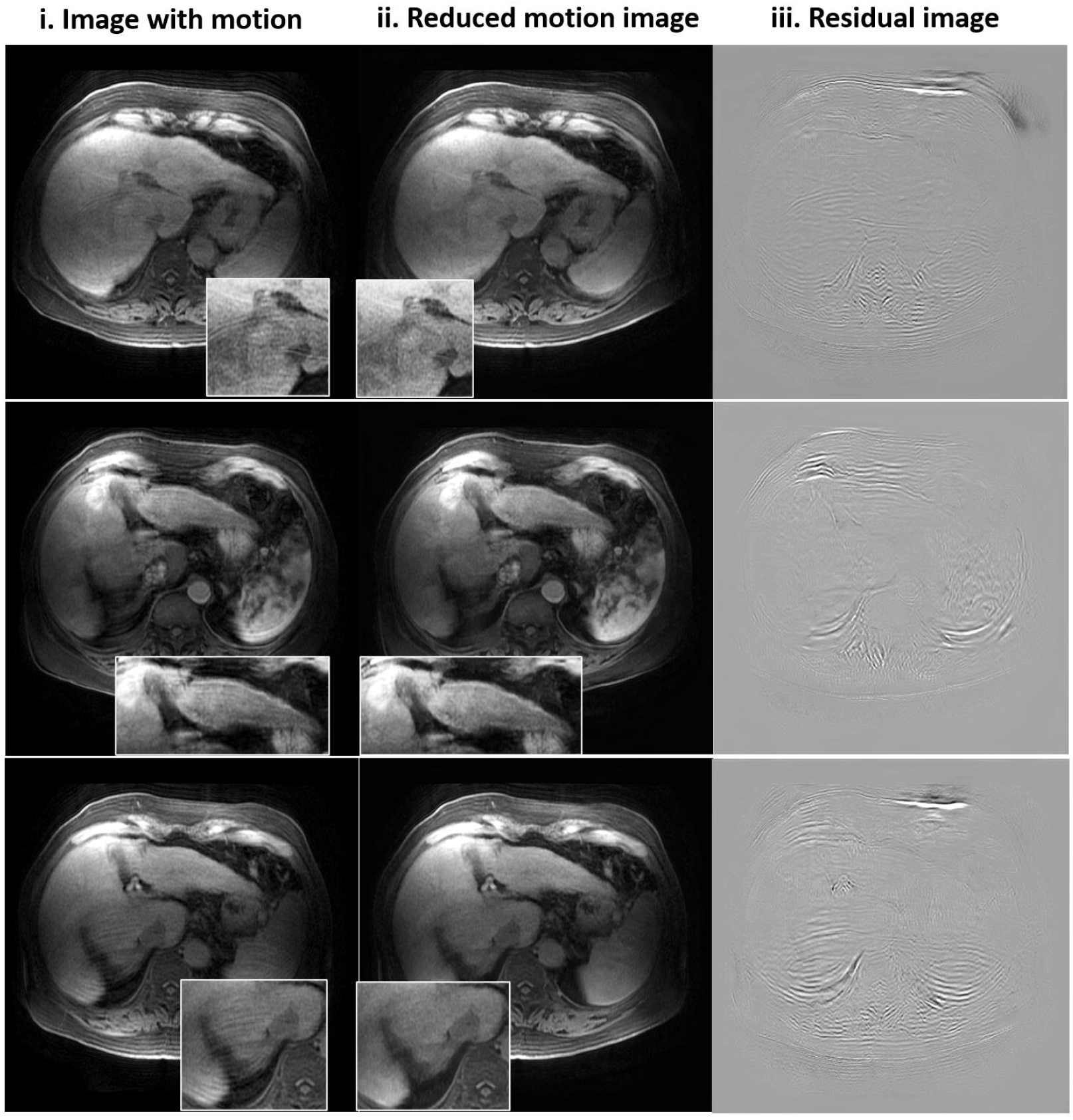
Motion-reduced images (column ii) generated by the J-net model from three testing patient images with real motion artifacts (column i). The residual images (column iii) were generated by subtracting the motion-reduced images (column ii) from the original images with real motion (column i).

## 4. Discussion

In this study, we developed a two-stage deep convolutional neural network, so called J-net model, to reduce motion artifacts on DCE-MRI of liver. The stage-I model was first trained to reduce the major artifacts of the images, followed by the stage-II model to restore the image details and prevent the over-smoothing problem. Our preliminary results demonstrated that this two-stage J-net model effectively improved DCE-MRI image quality with less motion artifacts.

It is more realistic to train the supervised DL model using simulated motion artifact images (16,18–20,24,25,36) because it is difficult to obtain image pairs of motion-corrupted images and corresponding repeated motion-free image. This is especially not possible for contrast enhanced acquisitions. Kromrey et al. added periodic phase errors to K-space lines to simulate periodic respiratory motion artifacts in abdomen MRI (20). Tamada et al. used random phase error patterns to simulate more-severe, non-periodic motion (19). Zaitsev et al. separated motion into three types: bulk motion, elastic motion, and respiratory motion (23). However, all previous studies only simulated one or two types of motion which may cause poor generalization of their motion correction models. Our study first simulated five different types of motion artifact mimicking real-life motion scenarios. We compared the model performance for these five types of motions. It showed relatively lower performance on sinusoidal motion compared with the other four types, indicating that the unpredictable nature (duration, frequency and phase) of the respiratory wave was more challenging for the model to correct, and this finding was in line with a previous study (37).

Only a few studies were focused on motion artifact reduction in DCE-MRI of the liver (19,20) and used the motion artifact reduction with convolutional (MARC) network with the pixel-by-pixel MSE as the loss function in their models. Although these methods achieved relatively good denoising effect on the liver MRIs, Yang et al. mentioned that using the pixel-by-pixel MSE as the loss function would lead to over-smoothing problems so that perceptually-important details were likely to be overlooked (30). Our unique two-stage network model aimed to solve this problem by reducing major motion artifact in stage-I and then recovering structural details in stage-II. The stage-I model adopted the architecture of DRN-DCMB network, which was previously used in brain MRI and led to the best performance of motion reduction among other state-of-the-art models (16). The DRN-DCMB network utilized multi-resolution blocks to extract motion details and meanwhile maintain desirable image contrast. The dense connections between the multi-resolution blocks compensated for information loss. In this study, using DRN-DCMB in the liver DCE-MRI cannot completely resolve the image over-smoothing issue. The same over-smoothing problem was observed in CT images and was believed due to the use of pixel-by-pixel MSE construction loss function (21). In order to solve this problem, the J-net model used the perceptual loss function that was computed from the feature maps with larger receptive fields, in a way similar to human visual perception for identifying objects (38, 39). Feature maps extracted from a deeper CNN layer (i.e., “block5_conv3” of the VGG-16 network in this study) that contained the most number of channels and higher resolution features can better preserve image details and structural information (26). Similar findings were reported to apply perceptual loss in deep CNN model to achieve artifact-free photon image reconstruction (40).

The performance of the J-net model (SSIM: 0.974 ± 0.016) outperformed the models reported in other studies of liver MRI motion reduction (SSIM: 0.91 ± 0.07) (19), even though different datasets were used. The unique design of the J-net model may also apply to retrospectively correct image artifacts on images of other body parts. The computational time of this model was fast for clinical use, taking 0.2 second to process and transform a motion-bearing image into a motion-reduced image.

There are several limitations of this study. First, clinical assessment of the model output images for interpretation of diseases and subtle anatomical structures was not performed. Second, the degree of motion reduction and image quality improvement was not evaluated by radiologists. Third, the effectiveness of motion reduction in real motion scenarios was inferior to that achieved in simulated motion scenarios. Future work includes increasing the size of training dataset with more comprehensive motion artifact simulations. Finally, other advanced networks such as generative adversarial network (GAN) (41) with its unique generator and discriminator can be exploited to generate super-resolution image to reduce image blurriness (42) and correct motion artifacts in MRI (43).

## 5. Conclusion

A two-stage convolutional neural network J-net model effectively reduced motion artifacts on liver DCE-MRI and overcame the image over-smoothing problem by integrating the perceptual loss function in the stage-II model. Once trained, this model offers the potential of improving the quality of abdominal MR images that are susceptible to motion in a real-time manner.

## Abbreviations

DCE-MRI: dynamic contrast enhanced magnetic resonance imaging
DRN-DCMB: deep residual network with densely connected multi-resolution block
CS: compressed sensing
CNNs: convolutional neural networks
DL: deep learning
3D: three-dimensional
FFT: Fast Fourier Transform
BN: batch normalization
LeakyReLU: Leaky rectified linear unit
MSE: mean square error
SSIM: structural similarity index
TR: repetition time
TE: echo time
FA: flip angle
MARC: motion artifact reduction with convolution

## Acknowledgements

This research is partially supported by the Swim Across America Grant from Rush Translational Sciences Consortium. The authors would also like to thank Dr. Sharon E. Byrd for her administrative support in establishing the medical imaging artificial intelligence research program.

## Notes

### Competing Interest Statement

The authors have declared no competing interest.

